# Structural characterization of human Vaccinia-Related Kinases (VRK) bound to small-molecule inhibitors identifies different P-loop conformations

**DOI:** 10.1101/112763

**Authors:** Rafael M. Couñago, Charles K. Allerston, Pavel Savitsky, Hatylas Azevedo, Paulo H. Godoi, Carrow I. Wells, Alessandra Mascarello, Fernando H. de Souza Gama, Katlin B. Massirer, William J. Zuercher, Cristiano R.W. Guimarães, Opher Gileadi

**Affiliations:** Structural Genomics Consortium, Universidade Estadual de Campinas — UNICAMP, Campinas, SP, Brazil; Centro de Biologia Molecular e Engenharia Genética, Universidade Estadual de Campinas, Campinas, SP, Brazil; Structural Genomics Consortium and Target Discovery Institute, Nuffield Department of Clinical Medicine, University of Oxford, UK; Aché Laboratórios Farmacêuticos SA, Guarulhos, SP, Brazil; Department of Biochemistry and Tissue Biology, Institute of Biology, State University of Campinas, Campinas, Brazil; Structural Genomics Consortium, UNC Eshelman School of Pharmacy, University of North Carolina at Chapel Hill, NC, USA

## Abstract

The human genome encodes two active Vaccinia-related protein kinases (VRK), VRK1 and VRK2. These proteins have been implicated in a number of cellular processes and linked to a variety of tumors. However, understanding the cellular role of VRKs and establishing their potential use as targets for therapeutic intervention has been limited by the lack of tool compounds that can specifically modulate the activity of these kinases in cells. Here we identified BI-D1870, a dihydropteridine inhibitor of RSK kinases, as a promising starting point for the development of chemical probes targeting the active VRKs. We solved co-crystal structures of both VRK1 and VRK2 bound to BI-D1870 and of VRK1 bound to two broad-spectrum inhibitors. These structures revealed that both VRKs can adopt a P-loop folded conformation, which is stabilized by different mechanisms on each protein. Based on these structures, we suggest modifications to the dihydropteridine scaffold that can be explored to produce potent and specific inhibitors towards VRK1 and VRK2.

## Introduction

Members of the Vaccinia-related kinases (VRK) family of serine/threonine protein kinases are present in the genomes of all metazoans and in those of poxviruses, including the family-founding member vaccinia virus B1R ^1-6^. The human genome encodes three VRK proteins. VRK1 is a nuclear kinase implicated in cell cycle control, chromatin condensation and transcription regulation, and its substrates include p53, Activating Transcription Factor 2 (ATF2), Activator Protein 1 transcription factor (c-Jun), Barrier to Autointegration Factor (BANF1) and histone H3 ^7-14^. VRK1 function is linked to cell proliferation and its overexpression has been associated with tumor growth ^14-17^. VRK2 is an active kinase that displays 2 alternative splicing forms, each of which localizes to distinct cellular compartments (cytoplasm and nucleus or ER and mitochondria) ^18^. The alternatively spliced C-terminal domain interacts with and regulates components of the JNK signal pathway (JIP-1, TAK1 and MKK7) and BHRF1, the BCL2 homolog in Epstein-Barr virus, independent of kinase activity ^19-21^. p53 and BANF1 are also substrates for VRK2^18,22^. VRK2 is also implicated in mitochondrial-mediated apoptosis^23^. The third VRK family member, VRK3, is not catalytically competent and is thus classified as a pseudokinase. VRK3 can bind and activate VHR, the phosphatase responsible for inhibiting the ERK signaling pathway ^8,10,24^.

The VRKs belong to the CK1 kinase group, whose members typically include additional structural elements within the conserved kinase fold. Crystal structures are available for the ligand-free kinase domains (KD) of VRK2 and VRK3 ^25^. A ligand-free, solution NMR structure is available for a C-terminal truncation of VRK1 containing the kinase domain and most of the regulatory C-terminal domain^26^. These structures revealed that all 3 human VRKs have the canonical kinase fold and possess a unique helix (αC4) between αC and β4. This helix links the two lobes of the kinase and is thought to maintain the VRK proteins in a closed conformation, characteristic of an activated state ^25^. VRK3 has a similar fold to VRK1 and VRK2 but displays a degraded ATP-binding site ^25^. The kinase domains of active human VRKs are similar to each other (~80% sequence identity) but only distantly related (<30% sequence identity) to those from other members of the CK1 kinase group.

In addition to the catalytic domain, VRK1 and VRK2 have large, non-catalytic C-terminal regions, which in VRK1 contains putative regulatory autophosphorylation sites ^26,27^. The solution structure of VRK1 revealed that this region interacts with residues from the protein ATP-binding pocket and activation segment ^26^. Ser/Thr residues within this region are phosphorylated ^10^, an event that may be necessary for the dissociation of the C-terminal domain from the ATP-binding pocket and activation of VRK1. Much less is known about the structure of the C-terminal domain of VRK2 and its impact on the kinase activity.

Here we present the first crystal structures for the kinase domain of VRK1 and the first crystal structures for ligand-bound VRK1 and VRK2. Our results reveal the structural changes necessary for the displacement of VRK1 C-terminal region by ATP-competitive inhibitors and suggest specificity determinants that may be employed to design small-molecule inhibitors selective for the two active human VRKs.

## Results

### Identification of potent VRK ligands

Previous studies using large libraries of diverse ATP-competitive inhibitors failed to identify potent hit compounds for VRK1 ^25,28^. To widen the scope of potential ligands, we analyzed previous results from thermal-shift assays (DSF) using VRK1_3-364_ and the published kinase inhibitor set (PKIS) ^29^. VRK2 was not included in the PKIS characterization study. For VRK1_3-364_, there were 29 compounds that displayed changes in melting temperatures, ΔTm, larger than 2.0°C (an arbitrary cut off for a positive hit in this experiment ^29^), with the top hit, GW297361X, displaying a ΔTm of 9.7°C (Fig. 1a; Supplementary Table S1). Compared to the other 67 kinases in the PKIS panel, VRK1 showed a relatively low number of hit compounds (Fig. 1a-b). Compounds displaying the highest ΔTms were quite promiscuous, as reflected by their low Gini coefficient (Fig. 1c). The Gini coefficient is a measure of compound selectivity, with values close to 1 representing highly selective compounds ^30^. The top hit GW297361X had a Gini coefficient of 0.4.

**Figure 1:**
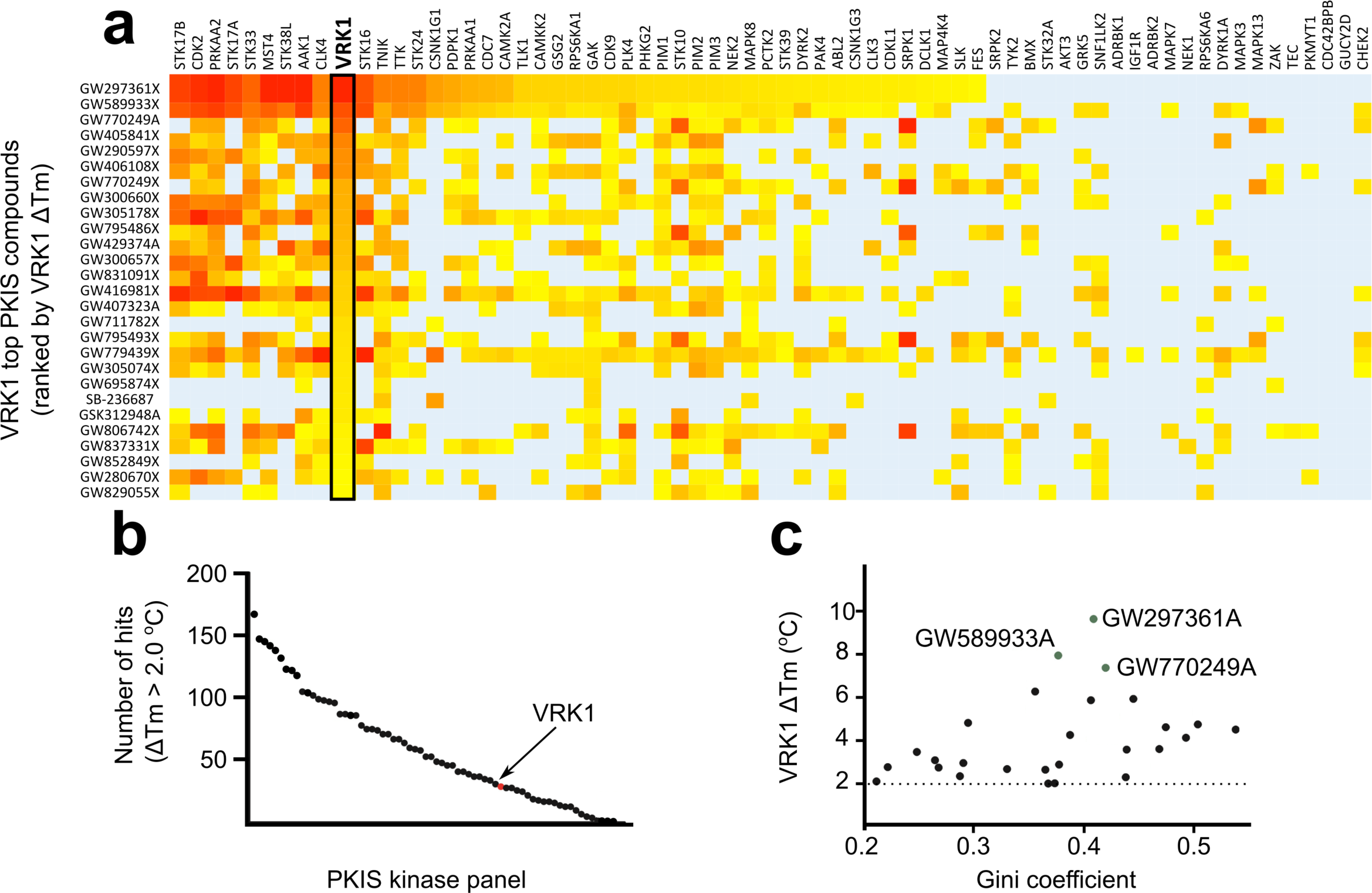
Analysis of published thermal shift assay (DSF) screening data for VRK1-PKIS. **(a)** Heat map showing DSF screening data for PKIS compounds with ΔTm > 2.0°C for VRK1 (black box) together with the results for other 67 kinases (x-axis). Compounds were ranked (top to bottom) according to decreasing ΔTm for VRK1. Background color indicates ΔTm as a gradient from 2°C (yellow) to the maximum observed ΔTm for particular proteins (red). Light blue background color indicates ΔTm < 2.0°C. (**b**) Graphic showing number of PKIS compounds with ΔTm > 2.0°C (y-axis) for VRK1 (red circle) and other 67 tested kinases (x-axis, black circles). (**c**) Graphic showing the poor relationship between ΔTm and Gini (selectivity) coefficients for PKIS compounds interacting with VRK1. The top 3 VRK1 hits are highlighted.

Within the PKIS, VRK1_3-364_ interacted mostly with broad spectrum compounds from an oxindole series derived from a medicinal chemistry program targeting CDK2 inhibitors ^31^. The PKIS data was also used in a hierarchical clustering (HCL) analysis to identify kinases with binding preferences similar to that of VRK1. These analyses revealed that the best VRK1 hit compounds also induced large ΔTm shifts on kinases such as CDK2 and TNIK (Supplementary Figure S1). A structure-activity relationship analysis of oxindoles within PKIS suggested the sulfur atom in GW297361X thiazole ring and polar substituents at the 6-position might be important for VRK1 interaction (Table 1).

**Table 1.**
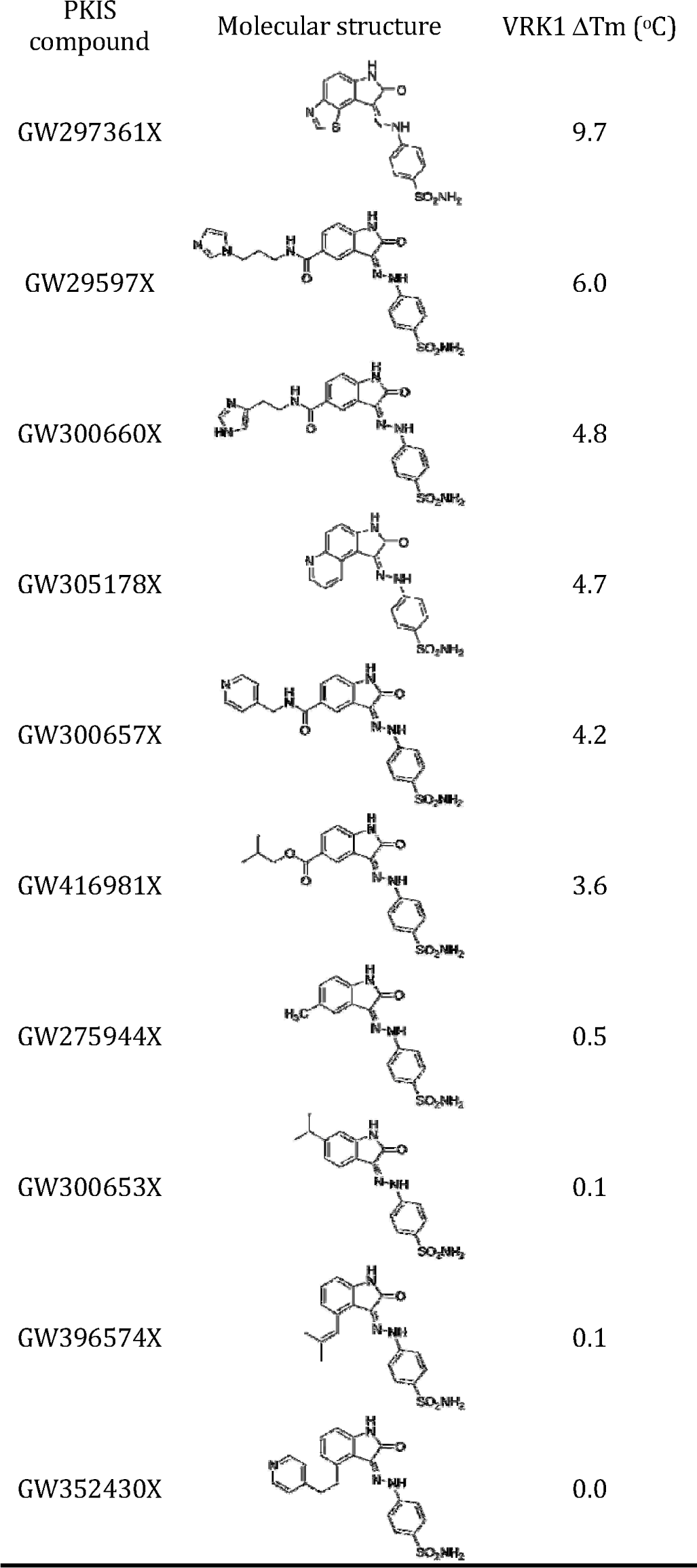
Structure-activity relationship of selected compounds from the oxindole series

|  |  |  |
| --- | --- | --- |
| PKIS compound | Molecular structure | VRK1 $\Delta T_m$ ($^{\circ}\text{C}$) |
| GW297361X |  | 9.7 |
| GW29597X |  | 6.0 |
| GW300660X |  | 4.8 |
| GW305178X |  | 4.7 |
| GW300657X |  | 4.2 |
| GW416981X |  | 3.6 |
| GW275944X |  | 0.5 |
| GW300653X |  | 0.1 |
| GW396574X |  | 0.1 |
| GW352430X |  | 0.0 |

Because the oxindole series in the PKIS showed poor kinase specificity, we sought to identify other potential inhibitors. Specifically, we performed a DSF screen on purified VRK1_3-364_ and VRK2_14-335_ using a commercially available library of 378 chemically diverse, bioactive, ATP-competitive, kinase inhibitors (Supplementary Table S2). To compare these results with those previously obtained for the PKIS compounds, we included GW297361X, the top VRK1 hit from PKIS, to the experiment. Note that the absolute ΔTm values for GW297361X differ between this and earlier experiments, possibly because of the use of different detection reagents and experimental setups.

The changes in melting temperature, ΔTm, for active VRKs spanned a wide range of values (Fig. 1). However, we found only a limited number of compounds with high ΔTm values for VRK1 and VRK2 (Fig. 2a-b and Supplementary Table 2) when compared to other protein kinases such as AAK1 (AP2-associated kinase 1; included for comparison)(Fig. 2c). For both VRK1_3-364_ and VRK2_14-335_, the compound inducing the largest ΔTm shift was BI-D1870, a dihydropteridinone inhibitor originally developed for p90 RSK (ribosomal S6 kinase) ^32^ which displayed larger temperature stabilization than the best PKIS hit for VRK1_3-364_, GW297361X (5.5 versus 3.9°C in this experiment, respectively). Within the identified compounds, four were able to induce ΔTm > 2.0°C for both VRKs, whereas three hits were exclusive to VRK1 and six hit compounds were exclusive to VRK2 (Fig. 4d). BI-D1870 was described as selective for RSK/N-terminal kinase domain and PLK1 (polo-like kinase 1) ^32^, but this report was based on a limited number of protein kinases. To understand the ligand binding site of active VRK proteins, we crystallized them in the presence of BI-D1870, the top DSF hit from the commercial library identified here. We also crystallized VRK1 with the top hit from PKIS GW297361X ^29^ and with broad spectrum kinase inhibitor ASC24 (compound **3** in ref ^33^). The three compounds used in crystallization are shown in Fig. 2e.

**Figure 2:**
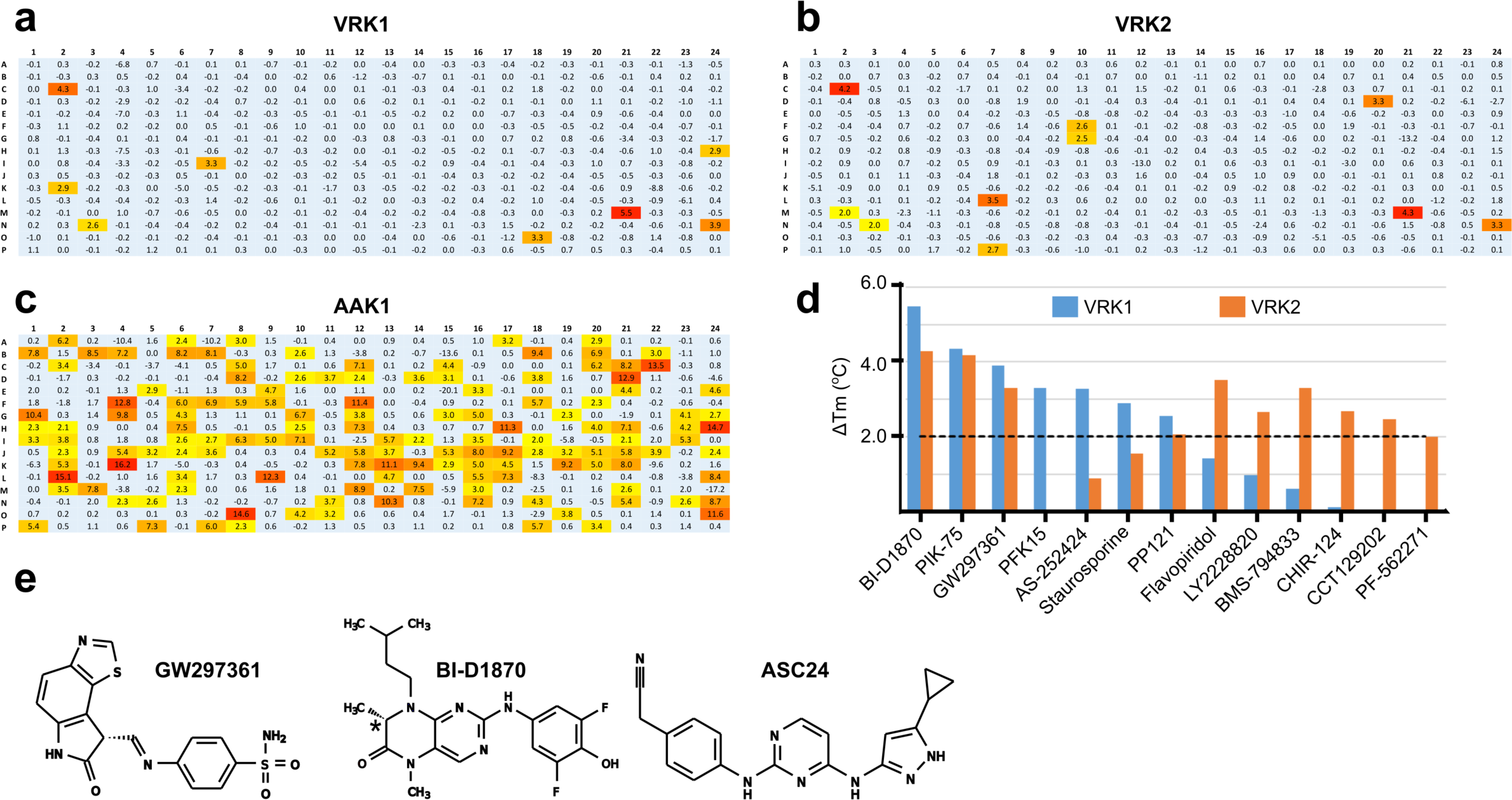
DSF screening identified few hit compounds for active VRKs. Heat maps showing DSF screening data for (**a**) VRK1, (**b**) VRK2 and (**c**) AAK1) included as comparison) versus a 378-compound library of cell-permeable, bioactive kinase inhibitors. The color scheme is the same as in Fig. 1a. Panels indicate rows (x-axis) and lines (y-axis) in a 384-well plate. Numbers indicate the observed ΔTm shift in degrees Celsius. (**d**) Bar graph showing ΔTm (°C) for top hit compounds for VRK1 (blue bars) and VRK2 (orange bars). (**e**) The three compounds used in crystallization. BI-D1870 was used as a racemic mix; the chiral carbon is marked with ^*^.

## Overall structure of ligand-bound active human VRKs

The ligand-free kinase domain of VRK2 has been crystallized previously ^25^. To crystallize VRK1, we mutated surface residues in order to reduce the entropic cost of crystallization ^34^. We obtained crystals of a construct containing the kinase domain and part of the regulatory C-terminal tail (residues 3-364) and 4 clusters of surface entropy reduction mutations (K34A/K35A/E36A; E212A/K214A/E215A; E292A/K293A/K295A and K359A/K360A)(Supplementary Figure S2). All four mutated clusters seem to contribute to crystallization as proteins mutated at only three of the clusters failed to crystallize. We then obtained co-crystal structures of VRK1_3-364_ bound to the three ligands - ASC24, GW297361X, and BI-D1870 (Fig. 2e); as well as of VRK2_14-335_ in the presence of BI-D1870. All structures were determined by molecular replacement and refined to resolutions between 2.0 and 2.4 Å (Table 2). For VRK1/ligand models, lack of electron density prevented model completion for some residues at the protein termini and for the glycine-rich P-loop in some polypeptide chains, likely because they are disordered in the crystal structures. The most complete VRK1_3-364_/ASC24 model contains residues 21-46 and 48-341 (chain A); and the one for the VRK1_3-364_/BI-D1870 complex contains residues 20-43, 49-341 (chain A). The VRK1_3-364_/GW92761X structure has a less ordered N-terminal domain and in two of the crystal polypeptide chains (C and D) models start after the protein P-loop region (residues 68 or 80). Nevertheless, the most complete polypeptide chain in this model contains an intact P-loop (chain A, residues 22-341; used for the structural comparisons below). The VRK2_14-335_/BI-D1870 model contains residues 14-328 and also has an intact P-loop.

**Table 2.**
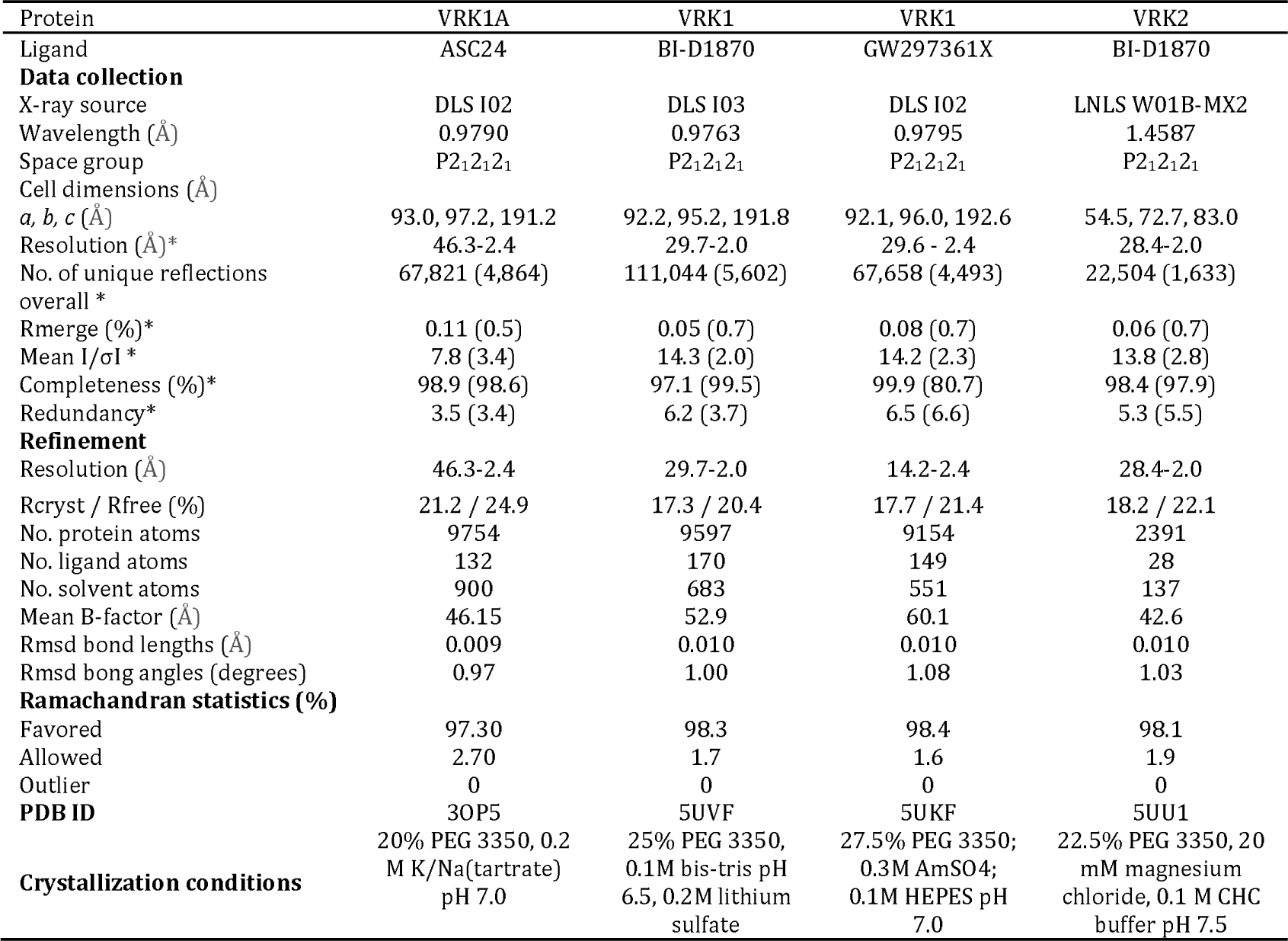
Crystallographic data

Figure 3 shows ligand-bound VRK1_3-364_ and VRK2_14-335_ proteins (Fig. 3a-d). All ligands bind to the ATP-binding pocket of the proteins. Ligand-bound VRK kinase domains adopt the canonical protein kinase fold, except for the presence of the additional α-helix between αC and β4 (αC4) exclusively found on VRK proteins. The equivalent region in VRKs’ closest structural homolog, casein kinase 1 (CK1) is a short loop ^35^ (Fig. 3e). VRK1_3-364_/ligand structures suggests the protein N-terminal region is quite dynamic, especially around the P-loop region (Fig. 3f). Structural comparisons of ligand-bound crystal structures and the structures of apo-VRK1 obtained by NMR in solution further suggest the protein N-terminal and P-loop regions are dynamic (Fig. 3g and Supplementary Figure S3).

**Figure 3:**
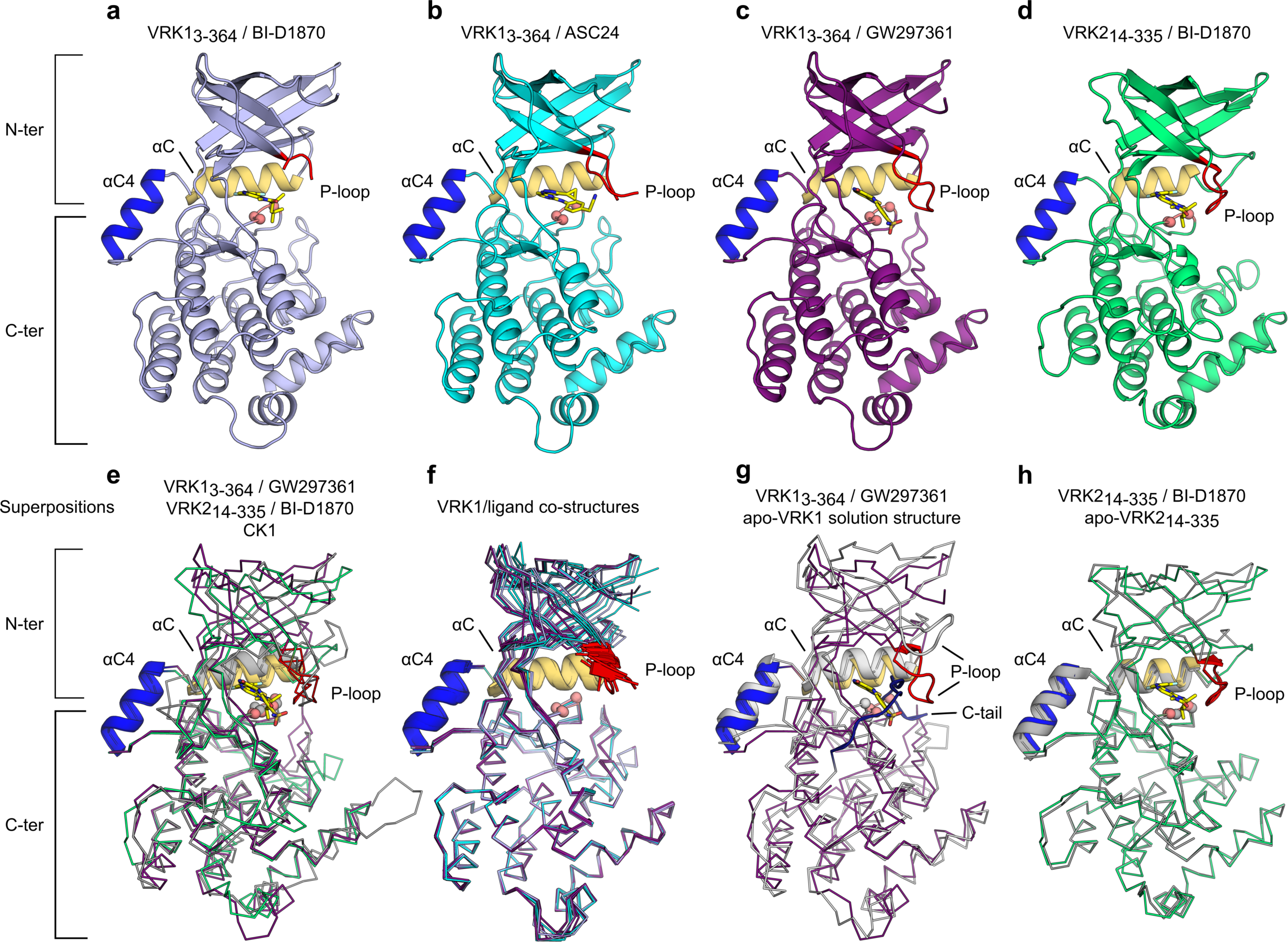
Co-crystal structures of ligand-bound active VRKs. Cartoon representation of VRK1_3-364_ bound to (**a**) BI-D1870, (**b**) ASC24 and (**c**) GW297361. (**d**) VRK2I_4-335_ in complex with BI-D1870. Position of αC (in yellow), DYG motif (Cαs shown as salmon spheres), αC4 (in blue) and P-loop (in red) are indicated. (**e**) Superposition of VRK1_3-364_/ GW297361, VRK2_14-335_/BI-D1870 and CK1 (PDB ID 1CKJ). Small molecule ligands are shown in stick model (carbon atoms in yellow; nitrogen atoms in blue, oxygen atoms in red and sulfur atoms in dark green). (**f**) Superposition of individual polypeptide chains in VRK1_3-364_/ligand co-crystals (colors as in panels **a-d**). (**g**) Superposition of VRK1_3-364_/GW297361 to the solution NMR structure (PDB ID 2LAV - in gray; residues 1-20 and 357-361 omitted for clarity). The position of the regulatory C-terminal tail (in dark blue) is indicated. (**h**) Superposition of apo- (in gray) and BI-D1870-bound VRK2_14-335_. In panels **e-h**, αC, αC4 and P-loop are show in cartoon representation, Cαs for residues within the DYG motif are shown as spheres and the rest of the polypeptide chain is shown as ribbons. Superpositions were performed using C-terminal residues only (residues 141-341 in VRK1).

Previous VRK structures revealed that αC4 plays an important role in stabilizing the closed, active conformation of the VRKs. In the active conformation, the kinase domain N- and C-terminal lobes are in close proximity to each other and a conserved Tyr residue from the DYG motif (DFG in most kinases) within the activation segment stabilizes a catalytically necessary ion pair between conserved lysine and glutamate residues ^25,26^. In most protein kinases, this closed conformation is only achieved following phosphorylation of regulatory sites in the activation segment. In the VRKs, however, these phosphorylation events are not necessary as αC4 promotes extensive contacts between N- and C-terminal lobes. The ligand-bound structures obtained here for active VRKs are in a closed, active conformation (Supplementary Figure S3) and display the same overall organization previously observed for the apo-structures, including the position of the VRK-exclusive αC4 (Fig. 3g-h). As for VRK1, changes between apo- and ligand-bound VRK2 structures are mostly located to the N-terminal and P-loop regions of the protein (Fig. 3h).

## Ligands can induce a folded P-loop conformation in the VRKs

The glycine-rich P-loop (GxGxF/YG motif) is located between the β-1 and β-2 strands and interacts with the phosphate groups in ATP, positioning the nucleotide for catalysis. In the absence of ATP, this region is mostly disordered due to the conformational flexibility conferred by glycine residues. The P-loop was disordered in the apo-VRK2 crystals and in most polypeptide chains within VRK1/ligand crystals. Kinases structures with an intact P-loop usually show this region in an extended conformation. To date, a folded P-loop has been observed in only a small number of kinase structures and has been associated with favorable inhibitor selectivity ^36^.

The crystal structures of VRK1_3-364_/GW297361X (chain A) and VRK2_14-335_/BI-D1870 reveal that the glycine-rich P-loop region of these kinases can adopt the so-called folded conformation. The folded conformation of the P-loop is stabilized by different types of interactions in VRK1 and VRK2 kinase domains (Fig. 4). In the VRK1_3-364_/GW297361X structure, the folded conformation of the P-loop is stabilized by a sulfur-π interaction between Phe48 and the sulfur atom in the thiazole moiety of GW297361X (Fig. 4a). Other than this interaction, VRK1_3-364_ P-loop residues do not contact atoms in the compound Fig. 4a-d). In VRK2_14-335_, both main chain and side chain atoms in the P-loop are in close proximity to atoms from the compound, whereas the side chain of the conserved aromatic residue in the P-loop (Phe40) could not be modeled due to lack of electron density - an indication that this residue is disordered (Fig. 4e-h). Thus, the folded conformation of the P-loop in VRK2 seems to be driven mostly by shape complementarity and hydrophobic contacts between the protein and the compound, whereas in VRK1 the P-loop folded conformation is held in place by a direct interaction between the compound sulfur atom and the protein aromatic residue. Moreover, the co-structure of VRK1_3-364_ bound to BI-D1870 revealed this compound cannot stabilize a P-loop folded conformation in this protein. These findings indicate the molecular bases for binding of small molecules and for stabilizing the P-loop folded conformation in VRK1 and VRK2 are different.

**Figure 4:**
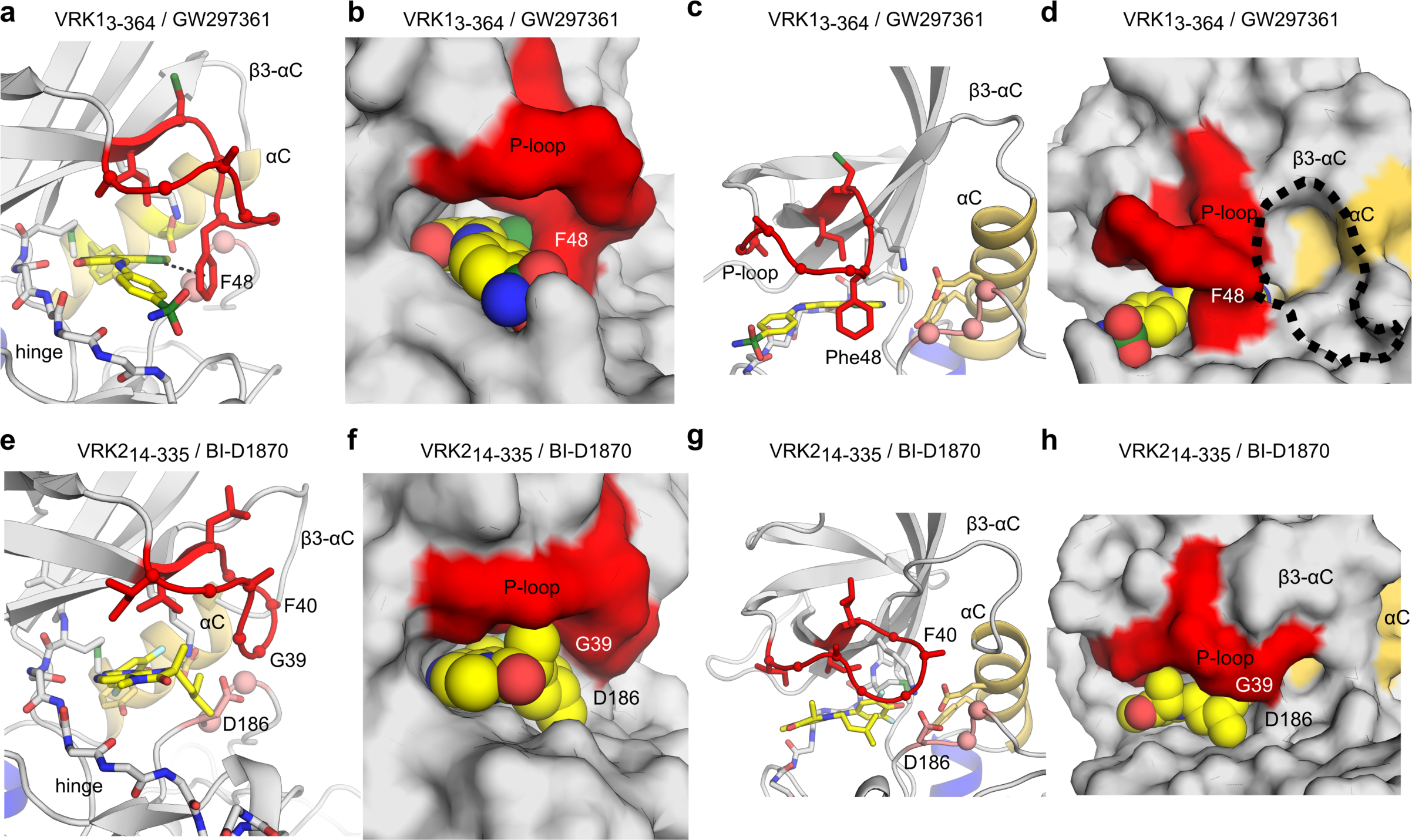
Ligand-induced, P-loop-folded conformation in active VRKs. **(a,c)** Cartoon and (**b,d**) surface representations of the P-loop region in VRK1_3-364_/GW297361 co-crystal. Dashed line in panel **a** indicates a π-π interaction between the sulfur atom in the thiozole ring from the ligand and P-loop residue Phe48. Dashed line in panel **d** indicates a cavity between αC and P-loop exclusive to VRK1. (**e, g**) Cartoon and (**f, h**) surface representations of the P-loop region in VRK2_14-335_/BI-D1870. Views in **a-b** and **e-f** are rotated ~90° from those in **c-d** and **g-h**, respectively.

Small molecule binding by VRK1 and VRK2 can also be distinguished by the precise positioning of the folded P-loop over the ATP-binding pocket. The folded P-loop of VRK2 points towards αC and fits under residues from loop β3-αC (Fig. 4g-h). By contrast, in VRK1, the folded P-loop points towards the solvent and the β3-αC loop points way from the P-loop, creating a small cavity between the P-loop and αC (Fig 4c-d). Thus, despite the high sequence similarity between the ATP-binding pockets of the VRK1 and VRK2 kinase domains, these structural rearrangements provide subtle differences that can be exploited during development of specific VRK inhibitors.

## Molecular bases for ligand specificity

Our aim was to identify a strategy to generate VRK1- and VRK2-selective inhibitors and our crystal structures reveal distinct structural features that may be explored to this end.

The binding of BI-D1870 reveals one approach that may be used to develop selective molecules for VRK2. BI-D1870 adopts a non-planar conformation that complements well the ATP-binding site of VRK2. BI-D1870 is synthesized as a racemic mixture. However, the VRK2/BI-D1870 co-structure revealed this protein favors a specific enantiomer of BI-D1870 (S-form) and that the methyl group attached to the stereogenic center projects into a cavity formed by the hydrophobic side chains from P-loop residues. The pendant isopentyl group in the bicyclic core is orthogonal to the plane of the ring in a configuration that allows Gly39 from the P-loop to approach Asp186 in the DYG motif, closing the P-loop over the compound 3,5-difluoro-4-hydroxyphenyl ring (Fig. 4e-h). Structural comparisons of ligand-bound and apo-VRK2 structures reveal the isopentyl moiety also orients the side chain of Asp186 towards the catalytic lysine. Further rearrangements in this region of the protein are induced by the compound 3,5-difluoro-4-hydroxyphenyl substituent - the gatekeeper methionine is displaced and the catalytic lysine re-positions itself to become the focal point in a network of polar contacts that also involves the phenol of BI-D1870 and the side chains of Asp186 and that from the conserved glutamate residue in αC, Glu73 (Fig. 5a). Residue Tyr77 in the αC helix also participates in this network via water-mediated hydrogen bonds to BI-D1870 and Glu73.

**Figure 5:**
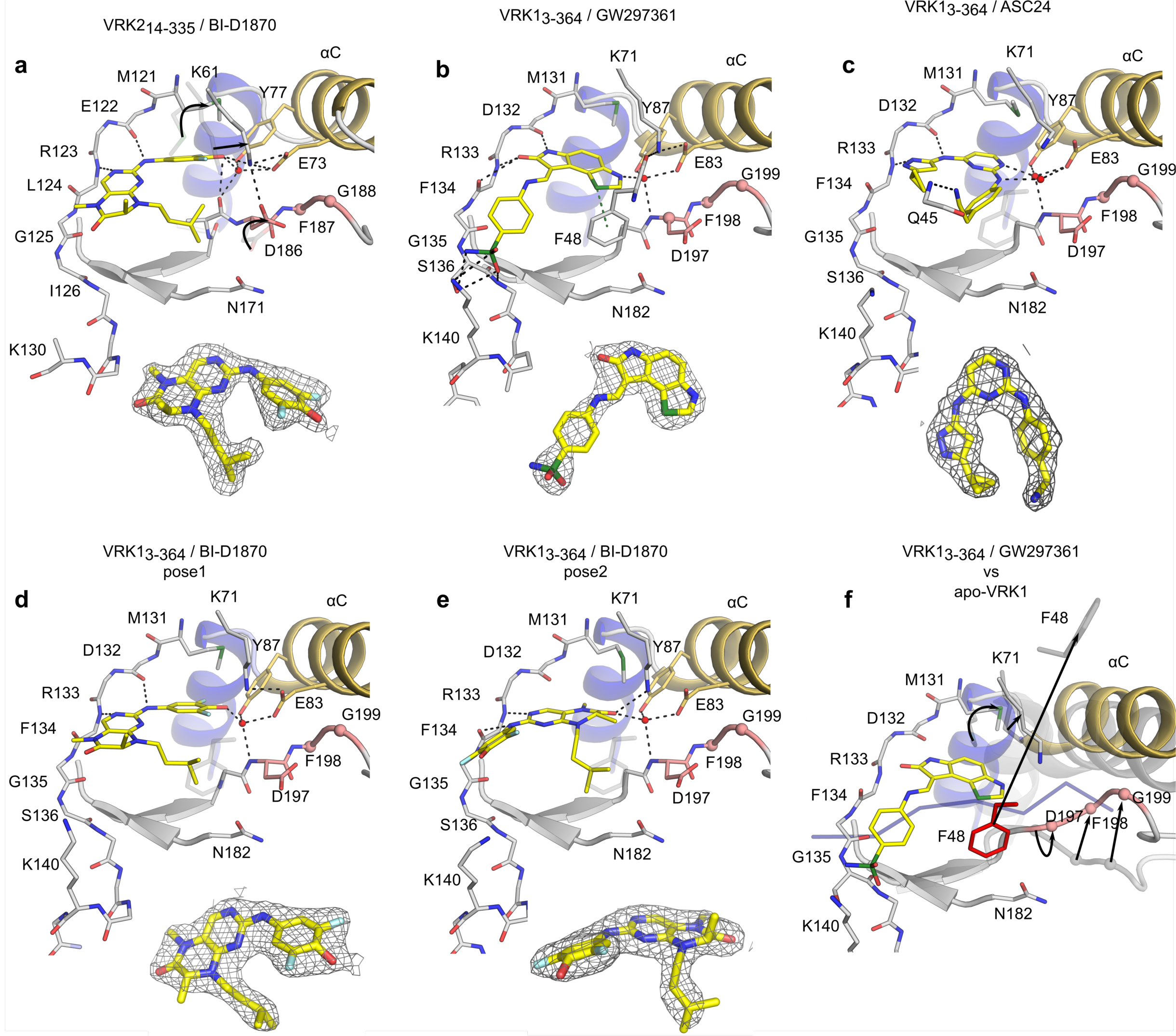
Crystallographic details of ligand binding. **(a)** Interactions between VRK2_14-335_ and BI-D1870. Arrows indicate re-arrangements of key residues compared to apo-VRK2 (PDB ID 2V62). (**b**) Interaction between VRK1_3-364_ and GW297361. The ligand sulfonamide group interacts with residues Ser136 and Lys140 at the C-terminal end of the hinge region. A green dashed line indicates a sulfur-π interaction between protein and ligand. (**c**) Interaction between VRK1_3-364_ and ASC24. The ligand acetonitrile group interacts with main chain amide from Gln45 in the protein P-loop. (**d-e**) Interaction between VRK1_3-364_ and BI-D1870. The ligand can adopt two distinct poses in VRK1 ATP-binding site. Electron density maps of the ligands (2Fo-Fc, contoured at 1.0 σ) are shown below each structural model. (**f**) Structural comparison of VRK1 ATP-binding site in complex with the regulatory C-terminal tail (NMR structure, PDB ID 2LAV, in faded gray; the backbone of the C-terminal peptide is shown in blue) and with GW297361. Arrows indicate rearrangements of key residues compared to the solution VRK1 NMR structure. Black dashed lines indicate potential hydrogen bonds. Red spheres indicate ordered water molecules. Cαs of residues within the DYG motif are shown as spheres. In panels **c-e** position of Phe48 from VRK1_3-364_ / GW297361 is shown for comparison (in faded gray). Color scheme as in Fig. 3.

Combined, the three VRK1 co-structures obtained here suggest strategies to develop selective molecules for this protein (Fig. 5b-f). These co-structures revealed VRK1 N-terminal domain to be quite dynamic, whereas the protein C-terminal lobe displays a more rigid architecture (Fig. 3f). Ligand GW297361 can stabilize the protein N-lobe via a sulfur-π interaction with P-loop residue Phe48. In this position, the side chain of Phe48 approaches those of Asp197 in the DYG motif and Asn182 at the ATP-binding site entrance to create a hydrophobic tunnel over the compound (Fig. 4a-b and e-f). Another notable interaction facilitated by GW297361 involves the sulfonamide group and residues located just outside the hinge region (Fig. 5b). The VRK1 binding pocket accommodates compound ASC24 in a tear-drop conformation and groups at opposite ends of the molecule reach towards residues in the protein P-loop. Despite these interactions, ASC24 does not support a P-loop folded conformation, most likely due to steric effects with P-loop residue Phe48 (Fig. 5c). Finally, BI-D1870 is seen in two different poses, supported by different hinge-binding modes. Both poses only loosely complement the volume of the ATP-binding pocket and neither pose supports the folded P-loop conformation, most likely due to steric effects with P-loop residue Phe48(Fig. 5d-e).

Structural comparisons of the solution NMR structure, where residues (346-354) within the C-terminal regulatory tail occupy the ATP-binding pocket, and the ligand-bound VRK1 structures reveal changes in the ATP-binding site are mostly limited to the protein DYG motif and to the P-loop. The position of the P-loop can vary widely depending on the identity of the ligand. On the other hand, amplitude of movement within the DFG motif is smaller. Binding of small molecule inhibitors to VRK1 repositions the catalytic lysine and facilitates formation of an ion pair with the conserved glutamate residue from αC, one of the hallmarks of the kinase active conformation (Fig. 5b and 5f and Supplementary Figure S2).

## Discussion

VRKs are implicated as mediators of a number of cellular processes and as potential therapeutic targets. But direct evidence for the roles of VRK1 and VRK2 in both disease and normal biology will be best derived from use of potent and selective pharmacological inhibitors of each active VRK ^37,38^. The crystal structures obtained here are the first example of ligand-bound VRK proteins. These structures expand our understanding on how these kinases interact with small-molecule ligands and reveal possible specificity determinants that can be explored by new inhibitors.

We found that the intrinsically active conformation of VRKs can be exploited to develop specific and potent compounds based on the dihydropteridinone scaffold. At 100 nM BI-D1870 was shown to inhibit all four RSK isoforms (1-4) (>98%) whilst also having activity against PLK1 (83%) and other 6 kinases (40-70%) in a 54-kinase selectivity panel, including CDK2 and CK1 ^32^. Our data show that BI-D1870 also targets the active VRKs and that this compound could serve as an advanced starting point for a specific inhibitor of these proteins.

The structures of RSK2/BI-D1870 (PDB:5D9K) ^39^ and of PLK bound to a similar compound (BI-D2536)(PDB 2RKU) ^40^ suggest that these proteins interact to the dihydropteridone chemotype in a dissimilar way to the VRKs (Fig. 6). Our structure of BI-D1870 bound to the constitutively-active conformation of the VRKs revealed that BI-D1870 binds to polar residues in αC that are not available in RSK2 and PLK1. We also found that the Tyr residue adjacent to the conserved glutamic acid in αC in the VRKs is substituted by aliphatic residues unable to form hydrogen bonds in most of the known BI-D1870 off-targets (notable exceptions include CK1, PLK1 and AurB, which have tyrosine, histidine or glutamine residues in this position). Thus, our data suggest that replacement of the hydroxyl group in the difluor ring of BI-D1870 with a strong hydrogen-bond acceptor may improve the interaction of the compound with the active VRK proteins and increase its specificity over the RSKs. Our analysis of the PKIS results also suggested the importance of a hydrogen-bond acceptor in this position. However, the energetic penalty for the potential water molecule displacement with larger substituents should be taken into consideration in order to increase potency.

**Figure 6:**
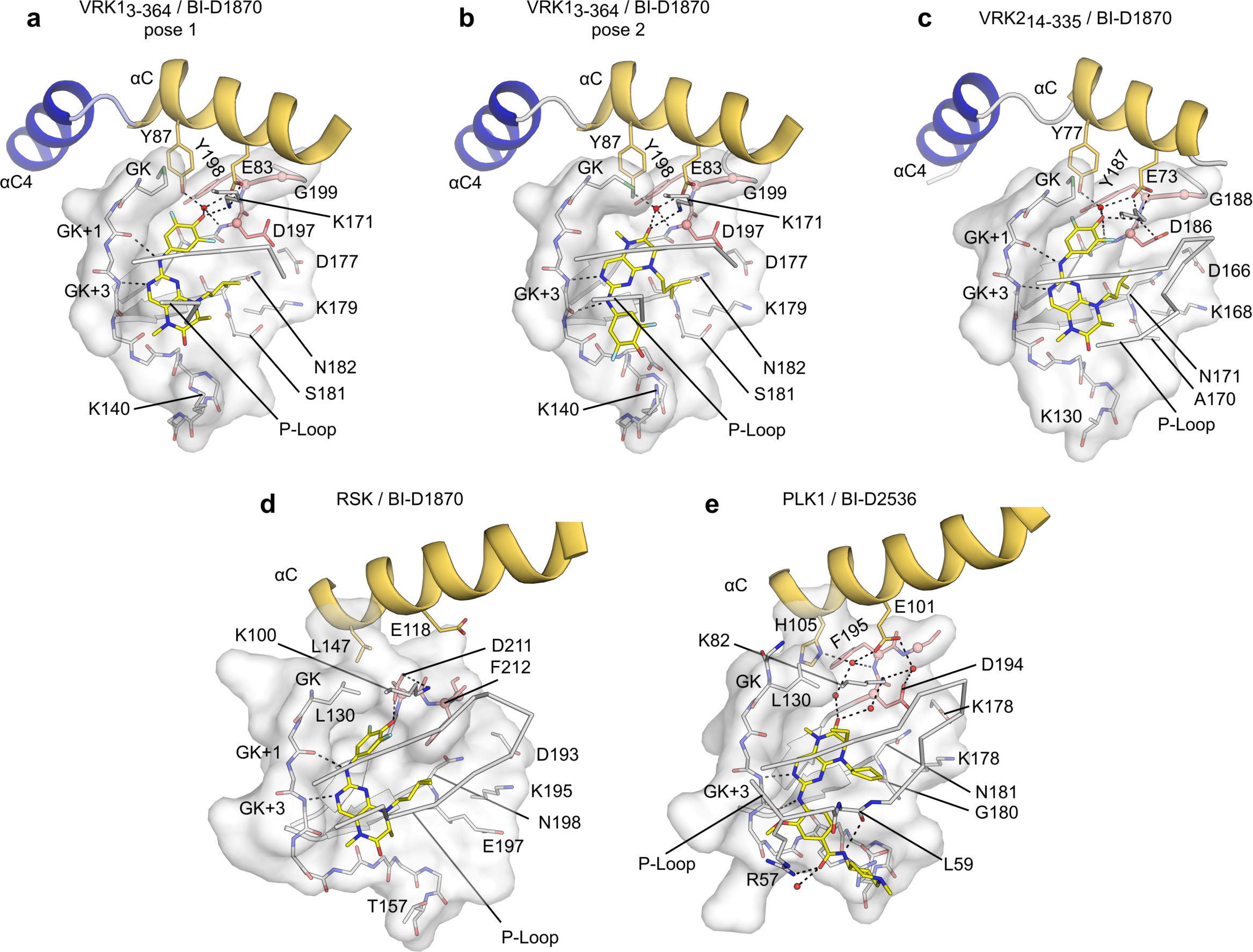
Active VRKs, PLK1 and RSK2 have dissimilar binding modes to pteridinone compounds. **(a-b)** BI-D1870 has two distinct binding modes to VRK1, supported by different interactions with the protein hinge region. (**c**) BI-D1870 interaction with VRK2. Both VRK1 and VRK2 are constitutively in a DFG-in, active conformation. (**d**) RSK2 binds to BI-D1870 in an inactive, DFG-out conformation (PDB:5DK9). (e) BI-D2536 binding mode to PLK1 (PDB: 2RKU) is similar to pose 2 in VRK1_3-364_/BI-D1870 co-crystals. PLK1 residues at the back of the ATP-binding site (K82, E101 and D194) contact the compound via an extensive network of water-mediated hydrogen bonds. Side chains of Asp and Tyr/Phe residues from the DFG (DYG in the VRKs) are shown in stick representations; Cαs in these residues are shown as spheres (salmon).

We also found that the stereogenic center in BI-D1870 might be explored to introduce specificity within the active VRKs. BI-D1870 is available as a racemic mixture but VRK2/BI-D1870 co-crystals favored a single enantiomer (S-form), in which the methyl group attached at the stereogenic center interacts with the protein P-loop. The co-structures of VRK1 bound to ASC24 and GW297361 suggest this protein can accommodate larger groups in this position. Thus, to generate a VRK1 selective series, we may extend or replace the methyl chain in the S-isomer. Likewise, extension of the methyl group in the R-form may prevent interaction with VRK2, while still allowing interactions with polar and charged groups at the binding pocket entrance in VRK1(Asp137, Gln139, Lys140, Asn182 Lys179, Asp137 - VRK1 numbering). The apparent preference of VRK2 for a single enantiomer may aid in the development of a negative control compound for a VRK selective chemical probe ^37^.

A promising feature of the ligand-bound VRK structures is the ability to adopt a folded conformation of the P-loop. This conformation has been associated with favorable inhibitor selectivity and has been observed for a number of protein kinases including major drug targets such as ABL1, Aurora A, FGFR1, cMET, p38 and MAP4K4 ^36,41^. The P-loop Tyr/Phe residue usually engages in aromatic stacking interactions, similar to the sulfur-π observed for VRK1/GW297361X structure. We propose that this interaction could be further explored to design a VRK1-specific inhibitor, as an equivalent interaction is unlikely to be supported by the folded conformation of VRK2 P-loop. For example, extension of the difluorinated ring to incorporate a sulfur-containing group, akin to the thiazole moiety in GW297361 may increase selectivity for VRK1. The highly dynamic N-terminal domain of VRK1 may also allow inhibitors to be accommodated in the observed gap between the P-loop and αC, akin to the recently-described binding mechanism of Erk1/2 inhibitor SCH772984 ^42^.

Although BI-D1870 is a good starting point for compounds targeting the active VRKs, development of specific inhibitors will require the elimination of cross-reactivity not only to other protein kinases, such as CDK2 and TNIK - identified in our HCL analyses; but also to un-anticipated off-targets. BI-D1870 interacts with the bromodomain of BRD4 with nanomolar affinity ^43^. Thus, future compound development strategies will also incorporate selectivity features identified from the structures of VRK1 bound to the promiscuous compounds ASC24 and GW297361, such as the P-loop interaction of ASC24 acetonitrile moiety and the ability of this protein to accommodate compounds in a tear-drop conformation.

## Methods

### Cloning, Expression, Purification and Crystallization

VRK1_3-364_ and VRK2_14-335_, appended with a tobacco etch virus (TEV) protease cleavable, N-terminal 6xHis tag, were expressed from vector pNIC28-Bsa4 ^44^. To improve VRK1 crystallizability, 4 clusters of surface entropy reduction mutations (SER) ^34,45^ were engineered into this protein- K34A/K35A/E36A; E212A/K214A/E215A; E292A/K293A/K295A and K359A/K360A. For protein production, BL21(DE3)-R3 cells, including a plasmid expressing lambda phosphatase ^46^, were cultivated in TB medium (supplemented with 50 μg.ml^-1^ kanamycin, 35 μg.ml^-1^ chloramphenicol) at 37°C until OD_600_ reached ~3 and then cooled to 18°C for 1 hour. Isopropyl 1-thio-D-galactopyranoside (IPTG) was added to 0.1 mM, and growth continued at 18°C overnight. Cells were collected by centrifugation and pellets suspended in 2x lysis buffer (lysis buffer is 50 mM HEPES buffer, pH 7.5, 0.5 M NaCl, 10 mM imidazole, 0.5 mM tris(2-carboxyethyl)phosphine [TCEP], Protease Inhibitors Cocktail Set VII - Calbiochem, 1/1000 dilution) prior to flash-freezing in liquid nitrogen. After thawing, cells were lysed by sonication on ice. Proteins were purified using Ni-Sepharose resin (GE Healthcare) and eluted stepwise in binding buffer with 300 mM imidazole. Removal of hexahistidine tags was performed at 4°C overnight using recombinant TEV protease while dialyzing against 1 L of gel filtration buffer (25 mM HEPES, 500 mM NaCl, 0.5 mM TCEP, 5% [v/v] glycerol). Proteins were further purified by reverse affinity in Ni-Sepharose followed by gel filtration (Superdex 200 16/60, GE Healthcare). Protein in gel filtration buffer was concentrated to 14 mg.ml^-1^ (VRK1_3-364_) or 20 mg.ml^-1^ (VRK21_4-335_) using 30 kDa MWCO centrifugal concentrators (Millipore) at 4°C. Compounds in 100% DMSO were added to protein solutions at 3-fold molar excess and incubated on ice for approximately 30 minutes. This mixture was centrifuged at 14,000 rpm for 10 minutes at 4°C prior to setting up 150-nl volume sitting drops at three ratios of protein-inhibitor complex to reservoir solution (2:1, 1:1, or 1:2). Crystallization experiments were performed at 20°C. Crystals were cryoprotected in reservoir solution supplemented with 20-25% glycerol before flash-freezing in liquid nitrogen for data collection. Diffraction data were collected at the Diamond Light Source (DLS) or at the Laboratório Nacional de Luz Síncrotron (LNLS). The best-diffracting crystals grew under the conditions described in Table 2. Crystal optimization used Newman’s buffer system ^47^.

### Structure Solution and Refinement

Diffraction data were integrated using XDS ^48^ and scaled using AIMLESS from the CCP4 software suite ^49^. Molecular replacement (MR) for VRK1_3-364_/ASC24 was performed with Phaser ^50^ using an ensemble of the following proteins apo-VRK2 (PDB ID 2V62), apo-VRK3 (PDB ID 2JII) ^25^ and CSK1 (PDB ID1CKJ) ^35^. The VRK1_3-364_/ASC24 structure was used as MR search model for phasing the other two VRK/ligand datasets. The apo-VRK2 structure (2V62) ^25^ was used as MR search model for the VRK2/BI-D1870 dataset. Automated model building was performed with Buccanner ^51^ following density modification with Parrot ^52^. Automated refinement was performed in PHENIX ^52^. Coot ^53^ was used for manual model building and refinement. Structure validation was performed using MolProbity ^54^. Structure factors and coordinates have been deposited in the PDB (see Table 2).

### Differential scanning fluorimetry (DSF)

Thermal stabilization assays were performed as described ^28,55^. Purified VRK1_3-364_ and VRK2_14-335_ were screened against a library of 378 structurally diverse, cell permeable ATP-competitive kinase inhibitors purchased from Selleckchem (Houston, TX, USA; catalog No. L1200). DSF experiments were performed in a 384-well plate format. Each well contained 25 µL of 1 µM kinase in potassium phosphate buffer and the Protein Thermal Shift dye at the recommended concentration of 1:1000 (Applied Biosystems; the composition of the buffer and the dye solutions are not disclosed). Compounds (10 mM) in DMSO were added to 16 µM final concentration to complete a total assay volume of 25.8 µL (3.1% final DMSO). Plates were sealed using optically clear films and transferred to a QuantStudio 6 qPCR instrument (Applied Biosystems). Fluorescence intensity data were acquired in a temperature gradient from 25 to 95°C at a constant rate of 0.05°C/sec and protein melting temperatures were calculated based on a Boltzmann function fitting to experimental data, as implemented in the Protein Thermal Shift Software (Applied Biosystems). Protein in 3.1% DMSO was used as a reference.

## Acknowledgements

SGC, a registered charity (number 1097737) that receives funds from AbbVie, Bayer Pharma AG, Boehringer Ingelheim, the Canada Foundation for Innovation, Genome Canada, GlaxoSmithKline, Janssen, Lilly Canada, Merck & Co., the Novartis Research Foundation, the Ontario Ministry of Economic Development and Innovation, Pfizer, the São Paulo Research Foundation (FAPESP grant 13/50724-5), Takeda, EU/EFPIA Innovative Medicines Initiative (IMI) Joint Undertaking (ULTRA-DD grant 115766) and the Wellcome Trust (092809/Z/10/Z). We thank Diamond Light Source for access to beam line I02 (proposal number 12988-1) and I03 (proposal number 10619-74) and beam line scientists Dr. Juan Sanchez-Weatherby and Dr. Carina Lobley who assisted with data collection. We also thank LNLS for access to beam line MX2 and beam line scientists Dr. Luciano Candido and Dr. Alexandre Lo Bianco Dos Santos who assisted with data collection. We thank Dr. Ricardo A.M. Serafim for help in preparing the SAR figure and Dr. Aled M. Edwards for critical review of the manuscript.

## Author contributions

PS and OG performed and analyzed the molecular biology experiments. PHG and OG designed, performed and analyzed the DSF experiments. RMC and CKA designed and performed the crystallography and structural biology experiments. RMC, HA and CKA analyzed the protein structures. CIW and WJZ performed initial PKIS analyses. HA, AM, FHSG and CRWG performed the PKIS analyses including the HCL, SAR and Gini coefficient determination. OG, RMC and HA wrote the manuscript. KBM helped coordinate and design the project. All authors reviewed the manuscript.

## Additional information

Accession codes: PDB accession codes are 3OP5 (VRK1_3-364_ / ASC24), 5UU1 (VRK2_14-335_ / BI-D1870); 5UVF (VRK1_3-364_ / BI-D1870) and 5UKF (VRK1_3-364_ / GW297361).

